# Graphene-based micro-electrodes with reinforced interfaces and tunable porous structures for improved neural recordings

**DOI:** 10.1101/2024.07.18.603459

**Authors:** Miheng Dong, Junjun Yang, Fangzheng Zhen, Yu Du, Siyuan Ding, Aibing Yu, Ruiping Zou, Ling Qiu, Zhijun Guo, Harold A. Coleman, Helena C. Parkington, James B. Fallon, John S. Forsythe, Minsu Liu

## Abstract

Invasive neural electrodes prepared from materials with miniaturized geometrical size could improve the longevity of implants by reducing the chronic inflammatory response. Graphene-based microfibers with tunable porous structures have a large electrochemical surface area (ESA)/geometrical surface area (GSA) ratio that has been reported to possess low impedance and high charge injection capacity (CIC), yet the control of the porous structure remains to be fully investigated. In this study, we introduce wet-spun graphene-based electrodes with pores tuned by sucrose concentrations in the coagulation bath. The electrochemical properties of thermally reduced rGO were optimized by adjusting the ratio of rGO to sucrose, resulting in significantly lower impedance, higher CIC, and higher charge storage capacity (CSC) than platinum microwires. Tensile and insertion tests confirmed that optimized electrodes had sufficient strength to ensure a 100% insertion success rate with low angle shift, thus allowing precise implantation without the need for additional mechanical enhancement. Acute *in-vivo* recordings from the auditory cortex found low impedance benefits from the recorded amplitude of spikes, leading to an increase in the signal-to-noise ratio (SNR). *Ex-vivo* recordings from hippocampal brain slices demonstrate that it is possible to record and/or stimulate with graphene-based electrodes with good fidelity compared with conventional electrodes.

## Introduction

Invasive neural electrodes are widely used in approaches for medical intervention and scientific research.^1–8^ Their proximity to neuronal cells can achieve high spatial-temporal resolution and SNR in the recording of local field potentials (LFPs), multi-unit activity, and single spike action potentials.^9^ The number of electrochemically active sites of neural electrodes is critical to the recording and stimulating capability. In general, invasive electrodes are designed with small geometrical sizes to minimize acute and chronic inflammatory response^10,11^ and to achieve high spatial resolution.^12–14^ Conventional invasive electrodes are generally based on metal substrates due to their outstanding electrical conductivity and mechanical properties.^15–18^ However, the smooth metal surface in conventional electrodes has a low ESA/GSA ratio, which limits performance. Additionally, corrosion of metal in physiological environments, results in the release of toxic metal ions and degradation of the electrode materials.^25–28^ This can result in inflammation and damage to surrounding tissues and reduce the reliability of electrode recordings. Therefore, increasing the surface area and minimizing corrosion are highly desirable in the design of neural electrodes.

Various methods have been employed to achieve these goals. Laser patterning, one such method to enhance electrochemical activity, only moderately improves performance.^19^ Surface coating using PEDOT has also been reported, but the integrity of the coating and the adhesion between metal and coating materials remains to be explored.^20,21^ In contrast, graphene materials have high corrosion resistance and biocompatibility.^29^ Moreover, the large surface area of graphene provides a high density of active sites for recording and stimulating neural signals, making it an ideal material for neural electrodes.^24^ However, there remains a poor understanding of how to control pore formation to fine-tune the graphene neural electrode properties, mechanical strength, and biocompatibility. To address this challenge, Aboutalebi *et al.* developed an optimized, commercially viable wet-spinning method.^30^ By exploiting the ability of acetone coagulant to extract water from the fibers rapidly, this method enables the fabrication of highly porous structures. Compared to other methods such as laser ablation^31,32^ and ice-templating,^33–35^ this approach offers finer control over pore formation, thus allowing for the optimization of neural electrode performance. However, without support from stiff substrates, the successful insertion of non-metal free-standing electrodes is a significant challenge as they do not have sufficient strength to penetrate brain tissue.^31^

In this study, we developed a strategy for fabricating graphene-based freestanding electrodes featuring tunable porous structures, which exhibit exceptional electrochemical and mechanical properties. This method involves the coagulation of graphene oxide (GO) dispersion to form hydrogel fibers with diffused sucrose incorporated within the structure. In the following thermal treatment process, sucrose is caramelized and carbonized, resulting in the formation of cavities and the release of gas, which creates pores while simultaneously reducing GO. Manipulating the sucrose concentrations in hydrogel fibers effectively fine-tunes the final porous structure. This structure was optimized by assessing impedance, charge injection capacity (CIC), and charge storage capacity (CSC). The low impedance was also confirmed through acute *in vivo* recordings. The discrete graphene flakes, reinforced by carbonized sucrose, demonstrated dramatically improved mechanical properties and longevity. In simulations of the surgical insertion process, optimized graphene-based electrode compositions exhibited a 100% insertion success rate with no change in morphology after an equivalent of 1.6 years of accelerated aging. Therefore, graphene-based electrodes are a potential candidate for *in vivo* recording and stimulation applications. This study provides insight into the mechanism of controlled pore formation in freestanding graphene-based microfiber electrodes and the potential for improving the performance of non-metal neural electrodes.

## Results and discussion

### Preparation and mechanism of graphene-based porous electrode

Graphene oxide (GO)/sucrose fibers were first prepared using a modified wet- spinning technique, as illustrated in Figure 1a. Initially, GO was dispersed in water to form a stable dispersion, facilitated by its functional groups. This dispersion was then extruded into a water coagulation bath containing CaCl_2_ and sucrose, forming hydrogel fibers. The coagulation process was driven by the electrostatic attraction between the negatively charged oxygen containing functional groups and the positively charged Ca^2+^ ions. Subsequently, the hydrogel fibers were dried to create dense GO/sucrose fibers. Thermal treatment was then applied to remove the functional groups from the GO flakes, restoring conductivity and creating porous structures (Figure 1b).^39,40^ The proportion of sucrose was regulated by adjusting its concentration in the coagulation bath used for diffusion. Dried GO/sucrose samples from the coagulation bath with sucrose concentrations of 0-50% were labeled GO-0 to GO-50, while the thermally treated samples were labeled rGO-0 to rGO-50. Figure 1c is the scanning electron microscopic (SEM) image of GO-50, and it is hypothesized that this fiber forms a porous GO framework with sucrose embedded in the pores, encased by a sucrose crust. It has been futher verified in figure 1d, showing the formation of pores when the heating temperature was increased to 120 °C, indicating that the crust could limit the shrinking process after dehydration at lower temperatures (90-120 °C),^41,42^ and this effect was also observed in GO fibers prepared with other sucrose concentrations (Figure S1). At higher temperatures (120-180 °C), sucrose melted, caramelized, and generated gas, forming voids within the fibers (Figures 1e, S2). A further temperature increase to 900 °C resulted in the reduction of GO and the carbonization of the caramelized sucrose, with no significant change in the cross-sectional geometry of the rGO-50 (GO-50 after thermal treatment), except for the formation of holes in the interior (Figures 1f, S3). In contrast, GO fibers prepared without sucrose (GO-0) underwent severe changes in geometry due to water evaporation and lattice changes, leading to irregular shrinkage with randomly distributed pores during thermal reduction (Figure S4).

**Figure 1.**
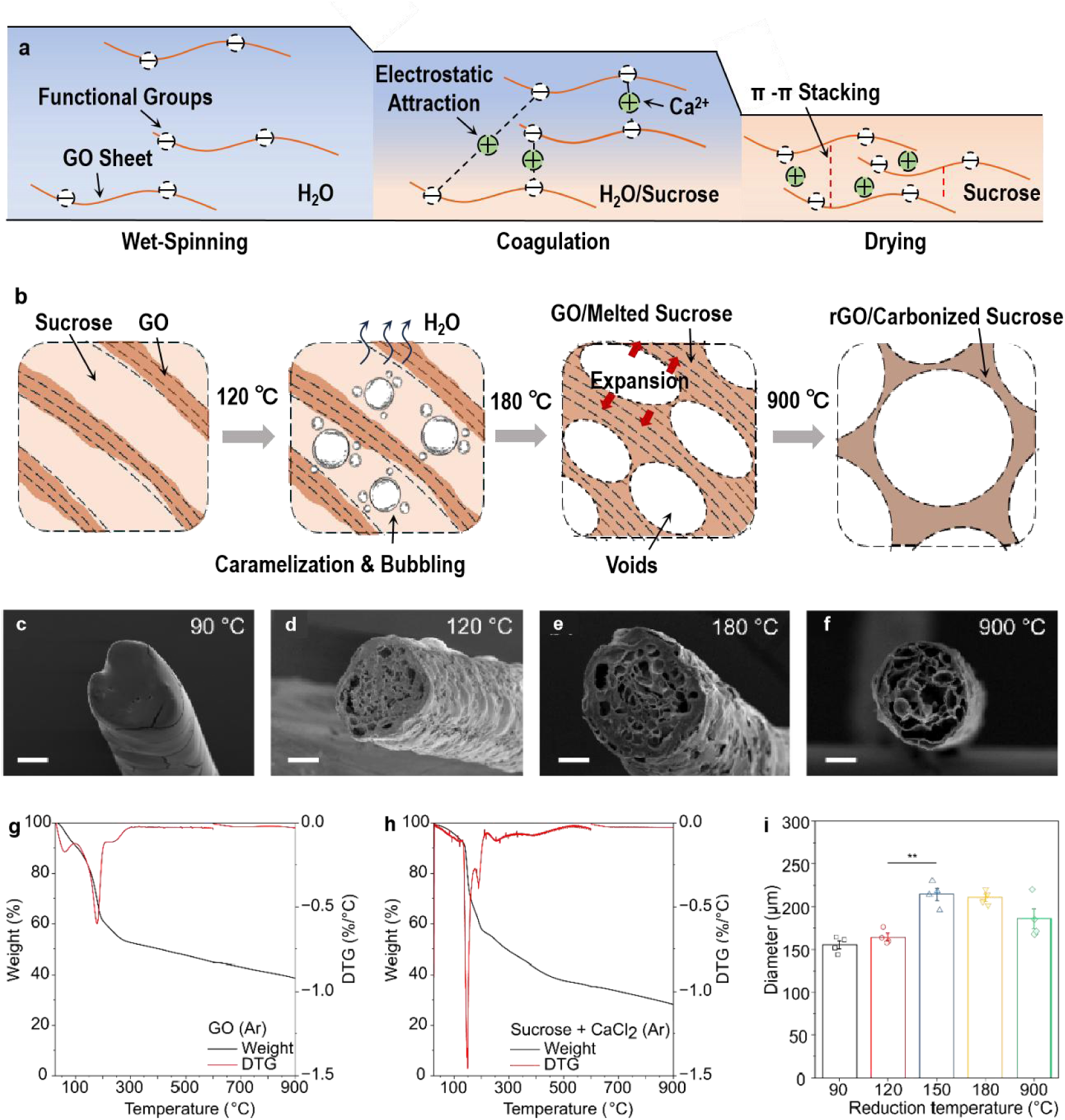
Mechanism of formation of porous rGO-based electrodes. Schematic illustrations of (a) the formation of GO/sucrose hydrogel fibers and (b) the mechanism of their evolution during thermal treatment. c) A GO fiber coagulated in bath 50 wt% sucrose (GO-50) exhibits a smooth sucrose shell after drying at 90 °C under vacuum. d-f) From 90–180 °C, released gas forms the porous structure of the GO-50 fiber. f) An rGO-50 fiber with a porous cross-section is prepared. g) The derivative of TG of GO is found with two broad peaks–the small peak below 100 °C (water evaporation) and a large peak (removal of water and GO functional groups). h) The derivative of TG of concentrated coagulation bath (sucrose+CaCl_2_) exhibits two peaks (caramelization of sucrose) below 200 °C. i) Diameter of GO-50 fibers at different reduction temperatures. Statistical analysis: Diameter data passed the Shapiro–Wilk test for normality (p > 0.05) and the Brown-Forsythe test for equal variances (p > 0.05). One-way ANOVA, n=4. Error bar: ± SEM. (****) p < 0.0001. Scale bars: c-f) 50 µm.

DTG analysis was performed to investigate the thermal behavior of GO and rGO samples. Two peaks at <100 °C and ∼180 °C were identified from the DTG graph of GO (Figure 1g). They were attributed to the evaporation of water and the reduction of GO, respectively.^43,44^ For the DTG graph of rGO-50, the rapid mass loss observed between 150 – 200 °C supports the caramelization process (Figure 1h), which involves the release of gas from the surface of GO fibers formed bubbles inside. The formation of gas also expanded the fiber and increased the diameter of the cross-section (Figure 1i). rGO-5 (fibers prepared with 5 wt% sucrose) was also found with the formation of pores between 120 °C and 180 °C and no significant change of the cross-section from 200 °C to 900 °C (Figure S5). The similarity in the critical temperature range of pore formation indicates that this mechanism is related to the properties of sucrose regardless of the concentration of sucrose used. To further clarify the effect of sucrose, GO-5 was soaked in 10 wt% CaCl_2_ to remove the sucrose in the hydrogel fiber. The fiber behaved very similar to GO- 0 between 90 °C and 150 °C in Figure S6. Irregular shrinkage and random voids were observed in cross-section, indicating the importance of sucrose in maintaining the geometry and generating pores.

In summary, the formation of the porous structure is driven by the caramelization process rather than the space occupied by sucrose. This claim is supported by two experiments. First, during the drying of the concentrated coagulation bath (sucrose & CaCl_2_), no clusters of sucrose crystals were observed, and the dried sucrose formed a solid, continuous structure (Figure S2a3). This suggests that in both the hydrogel and dried fibers, the sucrose was distributed homogeneously rather than in large clusters. Second, after the removal of iodine and sucrose from HI vapor-reduced dry GO-5 fibers, the cross-section did not exhibit pores like those observed in rGO-5 fibers, indicating that the dry GO-5 fibers did not contain voids occupied by sucrose crystals (Figure S7).

In addition to rGO-0, rGO-5 and rGO-50 fibers mentioned above, a range of graphene-based fibers were synthesized using various sucrose concentrations. From the cross-section of graphene-based fibers, it is clear that the pore size increases at higher sucrose concentrations (Figures 2a-d). The summary of porosity and fiber diameter for different groups is shown in Table S1. Among all groups, rGO-1 (88.7%) and rGO-5 (85.4%) exhibited the highest porosity (Figure 2e), while the porosity of graphene-based fibers in other groups was below 80%. The sucrose concentration has a broadly positive correlation with the fiber diameter of graphene-based electrodes, with higher sucrose concentrations tending to larger fiber diameters. The largest diameter was observed in rGO-50 (186.0 µm) (Figure 2f). This is attributed to the effectiveness of high sucrose concentrations in preventing fibers from shrinking during the drying process and releasing more gas in the following caramelization. The smallest diameter was found in rGO-1 (93.8 µm) and rGO-0 groups (103.9 µm). Since the cross-section of rGO-0 was not circular, the diameter of rGO-0 was calculated as the equivalent diameter of a circle covering the same area as the irregular cross-section.

**Figure 2.**
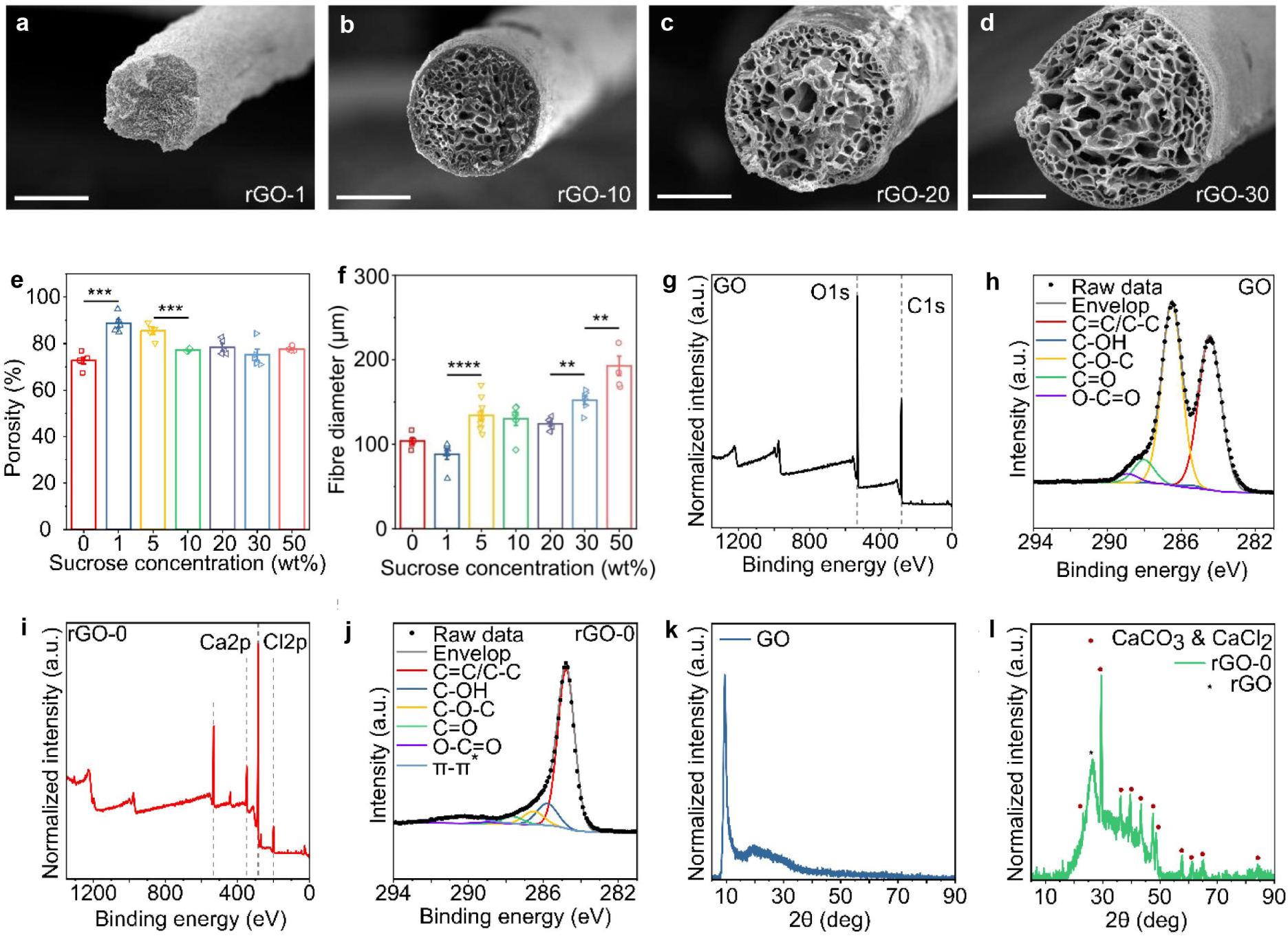
Geometry and characterizations of rGO fibers with different sucrose concentrations. Cross-sections of rGO fibers that reduced at 900 °C under vacuum. a-d) rGO fibers coagulated in baths with 1, 10, 20, and 30 wt% sucrose respectively. e) The porosity of different rGO fibers. f) The diameter of different rGO fibers. g, h) XPS survey and C1s spectra of GO. i, j) XPS survey and C1s spectra of rGO prepared with no sucrose (rGO-0). k, l) XRD spectra of GO and rGO-0, respectively. Statistical analysis: both porosity and fiber diameter data passed the Shapiro–Wilk test for normality (**p** > **0.05**) Fiber diameter data failed the Brown-Forsythe test for equal variances (**p** < **0.05**). One-way ANOVA was used for porosity data and Welch ANOVA was used for fiber diameter data. Error bar: ±SEM. (**) **p** < **0.01**, (***) **p** < **0.001** . Scale bars: a1-a4) 50 μm. n numbers in Table S1.

Thermal treatment can remove the chemical functional groups on GO flakes, thereby increasing the electrical properties of the fiber. For the application in neural prostheses, high conductivity is important for low-impedance recording.^36^ The functional group changes of GO fiber without sucrose were first investigated. XPS survey spectra revealed a clear oxygen peak in GO with an oxygen/carbon (O/C) ratio of 0.50 (Figure 2g). The C 1s spectrum (Figure 2h) confirmed that most of the oxygen existed in the form of ethers (46.7%, 286.5 eV), while carbon/oxygen bonds in other forms including C-OH (285.5 eV), C=O (288.0 eV) and O-C=O (289.0 eV) accounted for less than 10% (Figure 2h, Table S2).^45–47^ After thermal treatment (900 °C) under vacuum, the O/C ratio significantly reduced in both rGO-0 (0.17) and rGO-5 (0.13).^48, 49^ In rGO-0, there is a drop to 7.5% in C-O-C and an increase in C=C/C-C (74.9%) and C-OH (12.4%) ratio (Figures 2j, Table S3), demonstrating the reduction of GO.

A similar trend was also found in C 1s spectrum of rGO-5 which involved sucrose, where the C=C/C-C ratio increased to 63.1%, the C-OH ratio increased to 16.7% and the C-O-C ratio decreased to 10.8% (Figure S8, Table S4). Besides the carbon and oxygen peaks, calcium and chloride were also observed in rGO-0 and rGO-5 (Figures 2i and S8). This was caused by the use of CaCl_2_ in the coagulation bath, which remained during the drying process. In both rGO-0 and rGO-5, the major peak in C 1s became C=C/C-C bonds with significantly reduced intensity of ethers. Investigation of the thermal treatment for pure sucrose crystals revealed an O/C ratio of carbonized sucrose at 0.12 and a clear C-C peak in C1s spectra (Figure S9). Compared with pristine sucrose crystals, C 1s XPS analysis for the sucrose heated to 900 °C under vacuum revealed a clear decrease in the O/C ratio and a shift of the major peak from C-O to C-C, both of which indicate the carbonization of sucrose after thermal treatment.^50^ For XRD of GO samples (Figure 2k), the 2θ peak was located at 9.5°, indicating their oxidized state.^51, 52^ After thermal reduction, a broad 2θ peak of rGO appeared, centered at 26.4° (Figure 2l).^53^ Besides, the XRD spectra of pristine graphite powder exhibited a strong 2θ peak at 26.6° (Figure S10). These results from XPS and XRD collectively confirmed the effective reduction of GO after thermal treatment. It is also noted that CaCl_2_ and CaCO_3_ peaks also existed from the impurities of CaCl_2_ reagents in the reduced rGO/Ca sample.

### Electrochemical and electrical properties of rGO-sucrose fibers

To evaluate these properties of graphene-based fibers with different sucrose concentrations, a three-electrode system was employed. The graphene-based fibers served as the working electrode, a platinum wire served as the counter electrode and a Ag/AgCl electrode served as the reference electrode. The electrochemical measurements were performed in phosphate-buffered saline (PBS) solution at room temperature. The electrochemical impedance spectroscopy (EIS) measurements revealed that the impedance of all rGO groups from 0.1 Hz to 1,000 Hz was lower than that of Pt microwires (Figure 3a), indicating the potential to be used for recording slower neuronal responses (e.g. brainwave activity). Furthermore, the magnitude of impedance at 1 kHz confirmed differences between groups with statistical significance (*p* < 10^−2^^3^). In comparison to Pt microwires, lower impedance was found in all rGO groups from 0 to 50 wt%. In rGO groups with various sucrose concentrations, rGO-1 (0.49 Ω cm^2^), rGO-5 (0.57 Ω cm^2^) and rGO-10 (0.61 Ω cm^2^) exhibited lower impedance than rGO-0 (1.57 Ω cm^2^) from Figure 3e. From the phase plot, graphene-based electrodes exhibited more resistive behavior compared to the Pt microwires (Figure 3b). Additionally, there was a significant difference in CSC values among groups (*p* < 10^−9^) revealed by electrochemical characterization using cyclic voltammetry (CV). This can be visualized in the cyclic voltammetry curves of Pt and graphene-based fibers (Figure 3c). In the range of 0 to 20 wt% sucrose concentrations, a significant increase in CSC was measured for graphene-based fibers compared to pristine Pt microwires. Within rGO groups, rGO-5 exhibited the highest CSC value (404.55 mC cm^-2^) (*p* < 0.05), significantly outperforming the other groups (Figure 3f). Charge Injection Limit (CIL) characterizations exhibited a negative-leading bi-phasic current was increased until the voltage on the electrodes reached the water window of PBS (-0.6 V) (Figure 3d). There was also a significant difference between the groups (*p* < 10^−7^). Graphene-based fibers between 0 and 30 wt% sucrose groups were measured with higher CIL than Pt microwires (Figure 3g). Similar to the CSC results, rGO-5 (877.90 µC cm^-2^) was found with the highest CIL followed by rGO-1 (432.12 µC cm^-2^) (*p* < 0.01). No difference was found between rGO with 0, 1, and 10 wt% sucrose groups (*p* < 0.001). Results of impedance, CSC, and CIL are tabulated in Table S5. True conductivity equals the nominal conductivity divided by the volume percentage of materials in the fiber. There were significant differences between the true conductivity of rGO groups (*p* < 10^−8^).

**Figure 3.**
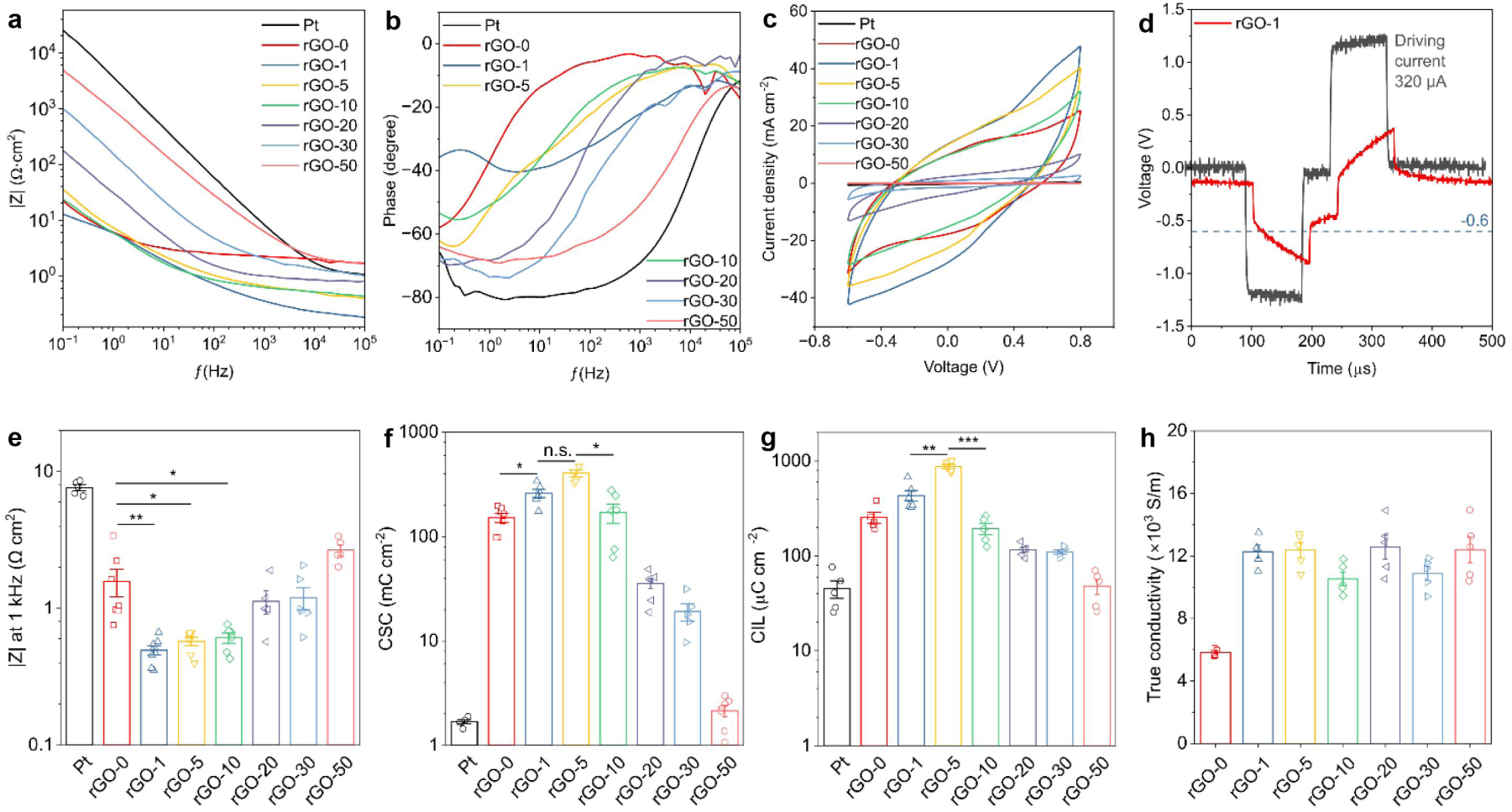
Electrical and electrochemical properties of graphene-based fibers with different sucrose concentrations. a) bode plot and b) phase plot of Pt and graphene-based fibers from 0.1 to 105 kHz. c) Cyclic voltammetry spectra of Pt and graphene-based fibers. d) An example of negative-leading bi-phasic pulse currents (320 µA, 100 µs/phase) driving an rGO-1 electrode to reach the lower limit of 1 × PBS water window (-0.6 V). Statistical analysis: each group of e) |Z| at 1 kHz, f) CSC, g) CIL, and h) true conductivity data passed the Shapiro–Wilk test for normality (*p* > 0.05). CSC and CIL data failed the Brown-Forsythe test for equal variances (*p* < 0.05). One-Way ANOVA was used for |Z| at 1 kHz and true conductivity and Welch ANOVA was used for CSC and CIL data. Error bar: ± SEM. (n.s.) not significant, (*) *p* < 0.05, (**) *p* < 0.01, (***) *p* < 0.001.

The results showed that the use of sucrose doubled the conductivity of graphene-based fibers (*p* < 10^−4^) but no difference was found between different sucrose concentrations (Figure 3h, Table S6). The involvement of sucrose significantly changed the electrochemical properties of graphene-based fibers.

The electrochemical and electrical characterizations further supported the effective reduction of GO flakes. Graphene-based fibers without any sucrose during fabrication showed significantly better performance than conventional Pt microwires. The porous structure in rGO is the main reason for this phenomenon. rGO-5 fibers exhibited the most superior properties in CSC, magnitude of impedance at 1 kHz and CIL, whose performance was significantly better than the no sucrose group and groups with other sucrose concentrations. It clearly shows the important relationship between pore size, porosity, and electrochemical performance. The relatively low concentration of sucrose generated a highly porous structure with small pores and preserved a regular cross- section, which may significantly increase the contact area between fiber tip and electrolytes.^31,54^ Additionally, the high conductivity of graphene-based fibers using sucrose might also have contributed to the significant improvement in electrochemical performance.

### Mechanical properties of graphene-based fibers

The high stiffness of metal and silicon makes it easy to insert conventional electrodes into tissues. In contrast, freestanding graphene-based electrodes without a metal support were evaluated by both standard mechanical testing for rod-shaped samples and customized experiments to simulate the actual implantation process of a neural electrode. Conventional rGO fibers suffer from weak interlayer bonding, especially in porous fibers. This study enhances mechanical properties by bonding rGO flakes with carbonized sucrose, as shown in Figure 4a. The three main reinforcements are: (i) interlayer bonding, (ii) edge bonding, and (iii) surface coating with carbonized sucrose. It is hypothesized that the mostly amorphous carbon structure of carbonized sucrose provides additional strength, repairing the discreate structure of the fibers in micro- nanoscale. In tensile testing, the ultimate strength and Young’s modulus were directly compared without correction for porosity differences, as the measured values represent the overall mechanical properties of graphene-based fibers. The Young’s modulus of graphene-based fibers with sucrose from 1 to 20 wt% was significantly higher than that of rGO-0 (Table S7). Among these groups, rGO-5 exhibited a Young’s modulus of 1510 MPa (Figure 4b), significantly better than rGO-1 (980 MPa, *p* < 0.001), rGO-10 (852 MPa, *p* < 0.0001), rGO-20 (8 MPa, *p* < 0.0001) and rGO-50 (2.3 MPa, *p* < 10^−7^).

**Figure 4.**
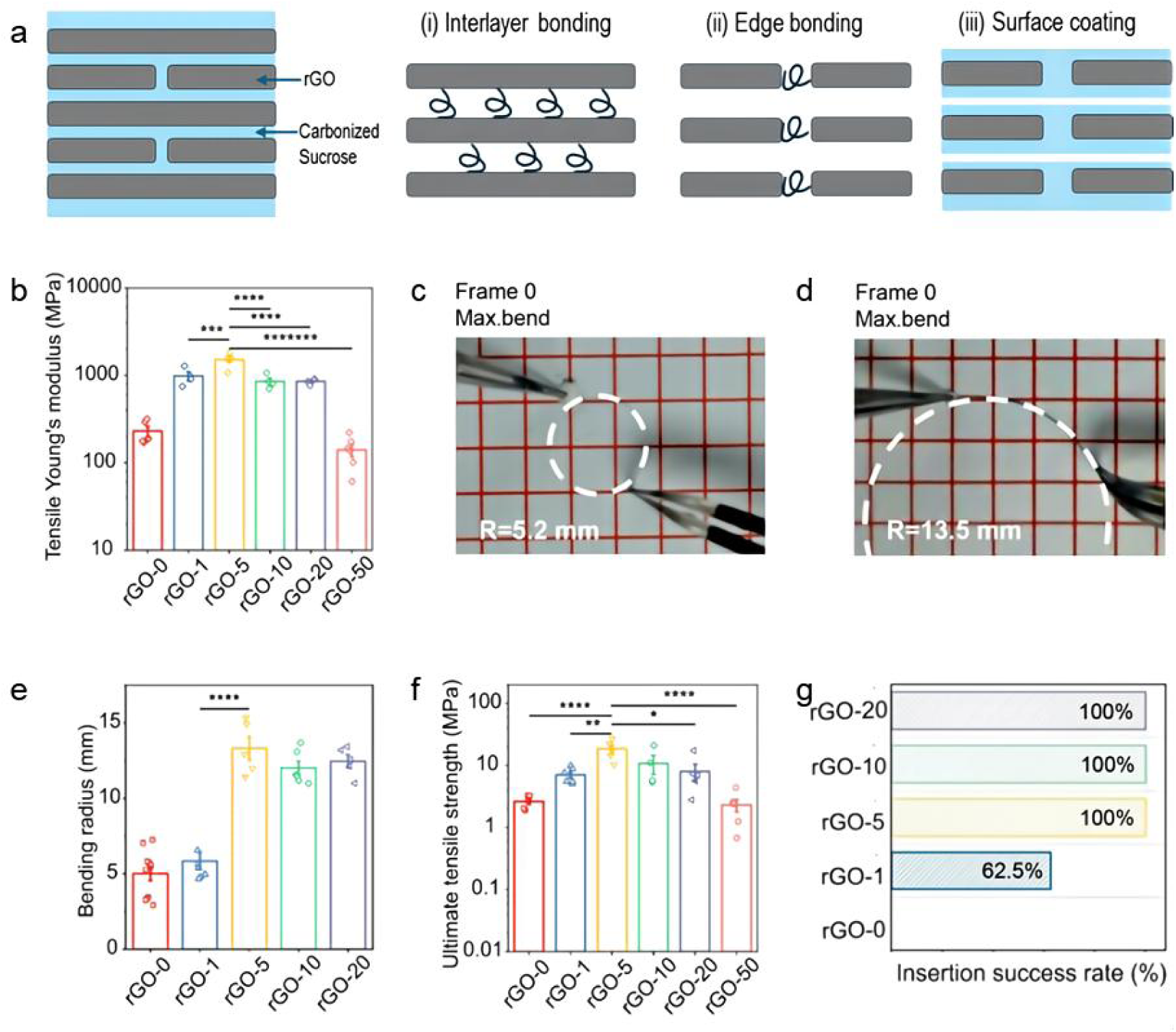
Mechanical properties of graphene-based fibers prepared from different sucrose concentrations. a) Schematic illustration of the mechanism behind the interlayer reinforcement of carbonized sucrose. b) Tensile Young’s modulus of graphene-based fibers. c) An rGO-0 fiber at maximum bend (R = 5.2 mm). d) An rGO-5 fiber at maximum bend (R = 13.5 mm). e) Bending radius of graphene-based fibers. f) Ultimate tensile strength. g) Insertion success rate for insertion test. Statistical analysis: b) Tensile Young’s modulus and f) ultimate tensile strength passed Shapiro–Wilk test for normality (***p*** > **0.05**) and the Brown-Forsythe test for equal variances (***p*** > **0.05**). One-Way ANOVA and Turkey post-hoc analysis were applied. b3) Bending radius data passed Shapiro–Wilk test for normality (***p*** > **0.05**) and the Brown-Forsythe test for equal variances (***p*** > **0.05**). One-Way ANOVA and Turkey post-hoc analysis were applied. d) Two-Sample t-Test. Error bar: ± SEM. (*) ***p*** < **0.05**, (**) ***p*** < **0.01**, (***) ***p*** < **0.001**, (****) ***p*** < **0.0001**, (*******)***p*** < **10^−7^**. Scale bars: c, d) the side of each red square is 5.0 mm.

Upon further strain of the fibers to the breakage point, rGO-5 with ultimate tensile stress of 18.3 MPa was also higher than rGO-0 (2.6 MPa, *p* < 0.0001), rGO-1 (7.1 MPa, *p* < 0.01), rGO-20 (8.0 MPa, *p* < 0.05) and rGO-50 (2.3 MPa, *p* < 0.0001) (Figure 4f, Table S8). Based on the results of Young’s modulus and ultimate tensile strength, it can be concluded that rGO-5 exhibited the best mechanical strength when the fiber is under tensile stress. The results indicate that a relatively small amount of carbonized sucrose can repair the discrete structure of the fibers. However, further increasing the amount may reduce mechanical strength due to the poor properties of amorphous carbon. It is also superior to fibers prepared by freeze-drying and chemical reduction by hydroiodic acid: Young’s modulus (350 MPa) and ultimate stress.^33^

During the surgery of implantation, electrodes are subjected to bending stress due to the manual handling of the microwire or the resistance of tissue during the insertion process. Manual handling by surgeons was simulated by holding a fiber at both ends and recording its shape change until it bent to break. For the rGO-0 fiber, the minimal radius of the fitted bending circle was 5.2 mm (Figure 4c) and the fiber fractured in the next frame (+33 ms) (Figure S11a). A rGO-5 fiber had a radius of 13.5 mm (Figure 4d) immediately before the breakage (Figure S11b). There was no significant difference in the radius of curvature between rGO-0 (5.0 mm) and rGO-1 (5.3 mm). Similarly, rGO-5 (13.3 mm), rGO-10 (12.0 mm), and rGO-20 (12.5 mm) also behaved similarly (Table S9). From the comparison between different rGO groups, it was concluded that the increase in sucrose concentration from 1 to 5 wt% leads to a significant increase in bending radius from 5.3 mm to 13.3 mm ((*p* < 0.0001) (Figure 4e).

The resistance a fiber experiences during the insertion process was simulated by inserting the fiber into a brain phantom made of agarose gel. The insertion speed was fixed by the customized stereotaxic apparatus. From the screenshots of the video footage, a graphene-based fiber would start to bend upon touching the surface of the gel. The rGO-0 fiber fractured when inserted into the phantom (Figure S12a) while a rGO-5 fiber successfully penetrated the gel (Figure S12b). The insertion success rate of rGO-0, rGO-1, rGO-5, rGO-10, rGO-20 are 0, 62.5%, 100%, 100% and 100% respectively (Figure 4f, *p* < 0.0001). This could be explained by the high ultimate strength and Young’s modulus of rGO-5 fibers. When a fiber is inserted, the angle shifts from the original direction, and the final direction is also calculated to evaluate the precision of implantation. Between rGO-1 and rGO-5 groups, the angle shift decreased from 3.3±0.4° to 0.9±0.2°. The high brittleness (small bending result in Figure 4d) of rGO-5 makes it unlikely to bend before penetrating the air-gel interface, thus significantly reducing the offset from the targeted position. No measurable angle shifts were recorded in rGO-10 and rGO-20. Results supported that 5wt% sucrose as the most effective in improving the mechanical performance of the freestanding graphene-based fiber in a simulated insertion process.

### Acute recordings and aging test of rGO-sucrose fibers

Electrophysiological recordings from the auditory cortex *in vivo* provide valuable insights into the neural processing of auditory stimulation. The quality of these recordings is influenced by the signal-to-noise ratio (SNR), which determines the clarity and interpretability of neural responses.^55^ Acute recording from the guinea pig auditory cortex was conducted to compare the signal-to-noise ratio of recorded response between graphene-based fibers and Pt microwires. The fabrication of recording electrode arrays follows a previously published protocol (Figure 5a).^47^ For each type of electrode, with the increase in sound stimulus (SPL), the SNRs between different SPLs are significantly different (Pt: p < 0.01, rGO-0: *p* < 0.0001, rGO-1: *p* < 0.0001, rGO-5: *p* < 0.001, rGO-20: *p* < 0.0001, One-Way ANOVA). SNRs recorded under 80 and 100 dB were higher than 60 dB because louder stimulus to the auditory cortex triggered stronger neuronal response (Figures 5b-d). The SNRs of all groups at different SPLs were tabulated in Table S10. At SPL 60 dB, SNRs of rGO-1 (9.6 dB) and rGO-5 (9.4 dB) were better than Pt (6.2 dB) (Figure 5b). Responses recorded from rGO-1 (14.9 dB) and rGO-5 (14.6 dB) at SPL80 dB were also found to be higher than Pt (10.1 dB) (Figure 5c).

**Figure 5.**
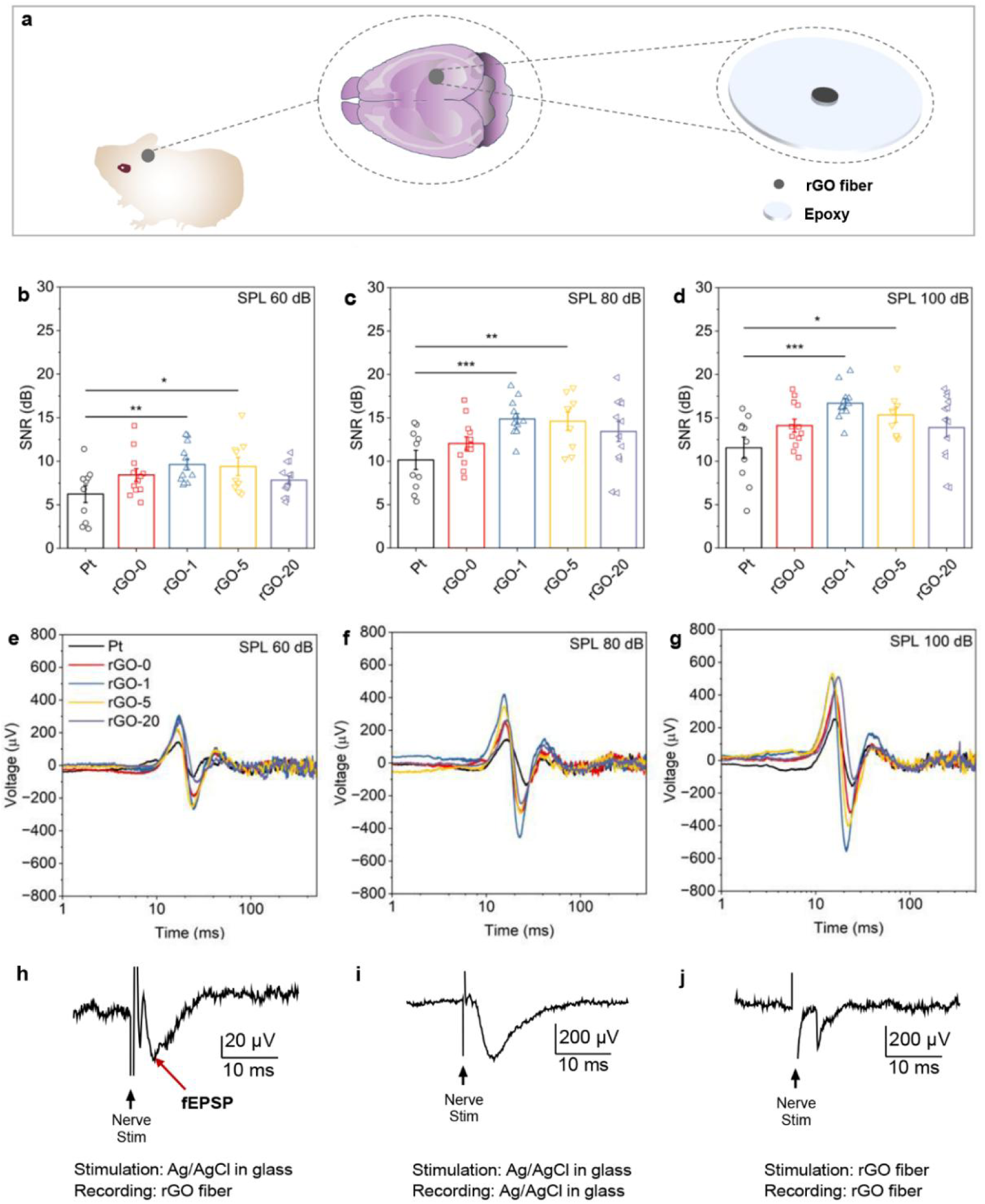
Evaluation of graphene-based electrode performance of recording. a) The manufacturing details of recording electrode arrays. SNRs of Pt and graphene-based fibers were recorded at b) SPL 60 dB, c) SPL 80 dB, and d) SPL 100 dB. Statistical analysis: graphene-based fibers compared against Pt using a Two-Sample T-test, n numbers in table S7. Error bar: ± SEM. (*) ***p*** < **0.05**, (**) ***p*** < **0.01**, (***) ***p*** < **0.001**. Recorded response of Pt and graphene-based fibers at e) SPL 60 dB, f) SPL 80 dB, and g) SPL 100 dB. h-j) Hippocampal brain slice recordings with various combination of stimulating and recording electrodes.

Similarly, at SPL of 100 dB, rGO-1 (16.7 dB) and rGO-5 (15.3 dB) also exhibited better recording capability than Pt (11.5 dB) (Figure 5d). At a fixed SPL loudness (60, 80, or 100 dB), rGO-1 and rGO-5 were two groups with higher SNR than Pt. This is explained by the high peak-to-peak amplitude of signals recorded from graphene-based fibers (Figures 5e-g).^56–58^ The above results demonstrate that graphene-based fibers offer superior SNR performance in acute auditory cortex recordings compared to traditional Pt microwires.

In addition, *ex vivo* hippocampal brain slice recordings and stimulations were performed using Ag/AgCl and graphene-based electrodes as stimulating and/or recording electrodes, respectively (Figures 5h-i). Ag/AgCl stimulating electrodes paired with Ag/AgCl recording electrodes produced the largest fEPSP (field excitatory postsynaptic potential). When used as a recording electrode, the RGO-50 electrodes recorded fEPSPs with a similar time course (panel h) to that recorded with an Ag/AgCl electrode (panel i) indicating good temporal fidelity of the rGO signal. The smaller size of the rGO-recorded fEPSP likely resulted from signal loss in the recording chamber due to the larger surface area exposed directly to the physiological bathing solution. Panel j shows that it is possible to both stimulate and record with graphene-based electrodes. The quicker response is due to the graphene-based electrode being much larger than the Ag/AgCl electrodes, therefore resulting in the activation of active responses/compound action potentials rather than the slower fEPSPs. Overall, graphene-based electrodes offer unique advantages for both recording and stimulation of neural activity, making them valuable tools for a wide range of electrophysiological research applications.

Carbonized sucrose forms amorphous carbon during the thermal treatment process. The electrodes were immersed in 0.9% saline (85 °C) for three weeks with constant charge-balanced stimulus to simulate the chronic implantation for 1.6 years.^36^ Through the comparison between cross-sections of randomly selected electrodes before and after accelerated aging, the porous surface of rGO-5 was well preserved (Figure S13). This indicates that the fibers with sucrose are potentially suitable for chronic applications.

## CONCLUSION

In this work, we unveiled the mechanism of pore generation in sucrose-reduced graphene oxide (rGO) fibers and present a novel approach to tailor their porous structure and electrochemical properties by modulating the sucrose concentration in the wet- spinning coagulation bath. This approach enables the fabrication of porous graphene- based electrodes (rGO-5) with excellent electrochemical performance, high surface area, and mechanical robustness. The promising potential of these electrodes was also demonstrated for neural interface applications, particularly in low-impedance, high signal-to-noise ratio in acute recordings of local field potentials and the ability to record fEPSPs in ex vivo brain slices. These findings provide new strategies for designing and fabricating graphene-based electrodes with tailored properties and advance the development of graphene-based electrodes for neural interfaces and other bioelectronic applications.

## MATERIALS AND METHODS

### Wet spinning and thermal reduction of GO fibers

GO dispersion at 19.1 mg g^-1^ (Hangzhou Gaoxi Technology Co. Ltd., China) was used as received. CaCl_2_ (≥96%, AR, Greagent) and sucrose (D-(+)-Sucrose, AR, Greagent) were dissolved at different concentrations in DI water. The solution was then centrifuged (Centrifuge 5430, Eppendorf) at 5,000 rcf for 3 min to remove insoluble particles. The supernatant was transferred into a glass petri dish as the coagulation bath. The petri dish was attached to a motor with a tuneable spinning speed. GO dispersion in a 5 ml syringe was injected in the rotating bath through a 25-gauge needle. The injection speed was controlled by a syringe pump (Model: LSP02-1B, LongerPump). The initial GO fibers prepared used a rotation bath (10 RPM). The needle was placed 70 – 80 mm away from the center of the bath. Fibers prepared from 2.5 and 5 mg g^-1^ GO coagulated in a 5 wt% CaCl_2_ bath were suspended vertically from one end to dry. Fibers prepared from the initial parameters were non-uniform. To achieve the best result, the final speed rotation bath was set at 70 RPM. After the GO dispersion coagulation into the GO hydrogel, the fibers were removed from the bath and suspended across two edges of a graphite crucible with both ends fixed to dry. The crucible with fibers was then placed in a tube furnace (OTF-1200X, MTI KJ Group) and reduced under vacuum (VRD-4, VALUE). The thermal reduction protocol is as follows: heat from room temperature to 600 °C at 1 °C min^-1^, heat from 600 °C to 900 °C at 5 °C min^-1^, kept at 900 °C for 1 hr and followed by natural cooling. For other reduction temperatures below 900 °C, the heating speed followed the same protocols. Cooled fibers were transferred onto silicone film for storage.

### Fabrication of graphene-based electrodes

The graphene-based fiber was connected to a copper leading wire using silver glue and fixed using a heat gun. The wire and fiber were then insulated using epoxy. Finally, a sharp blade was used to cut the insulated fiber to expose one end as the interface for electrochemical characterizations and neural recordings.

### Instrumental characterization of GO, rGO

For both x-ray diffraction (XRD, ARL EQUINOX 100 X-ray Diffractometer, ThermoFisher) and x-ray photoelectron spectroscopic (XPS, ESCALAB Xi+, ThermoFisher) characterizations, dried samples were ground into powder. Al Kα was used as an X-ray source for XPS.

### Electrochemical and electrical characterization of rGO

The potentiostat used is CHI660E (CH instruments, China). Platinum microwires (99.99% trace metals) as controls were used as received. The parameters of cyclic voltammetry and electrochemical impedance spectroscopy were similar to those used for ErGO-coated Pt/Ir electrodes. In the measurement of impedance, the driving sinusoidal voltage was 5 mV in amplitude and the scanning frequency was from 100,000 to 0.1 Hz. The measurement of charge injection limit has been found in previous publications from other groups.^36,37^ It was performed by applying a cathodic-first charge-balanced current pulse to the electrode using a programmable current source (KEITHELY 6221, Tektronix, US) and recording the voltage on the electrodes using a digital oscilloscope (TBS1000C, Tektronix, US). The current gradually increased until the maximum cathodal voltage (Emc) reached the lower limit of the water window (-0.6 V). To get the conductivity of fibers, a direct current was applied to a fiber using the current source, and the voltage across the fiber was recorded by a digital oscilloscope (TBS1000C, Tektronix, US). The diameter and area of the fiber tip were calculated based on optical microscopic images.

### Mechanical characterization of GO/sucrose and graphene-based fibers

Tensile stress tests were carried out using an electromechanical universal testing machine (TSC504B, China). In bending tests, two ends of the fiber were bonded to Blu Tack (Bostik, Australia). With one end fixed on the table, the other end was pressed with a tweezer to bend the fiber to fracture. The process was recorded using a video camera recording at 30 fps. The bending radius was calculated based on the frame before the fiber broke. In insertion tests, 0.6% agarose gel was prepared as described in the previous project of ErGO-coated Pt/Ir electrodes. The fiber was adhered to a plastic stick with Blu Tack. The stick fixed to the droplet dispensing equipment of a contact angle goniometer (SDC-200, SINDIN, China) was lowered down at a speed of 0.3 - 0.4 mm s^-1^. The video was captured with the camera on the goniometer.

### Acute recordings from guinea pig auditory cortex

Ethics approval for the a acute recording was obtained from St Vincent’s Hospital AEC, project #02/2. The surgery and recording followed a previously published protocol.^38^ Two guinea pigs were used, and the sampling rate was 100 kS s^-1^. In the comparison between graphene-based electrodes with different sucrose concentrations, the calculation of SNR used the peak-to-peak voltage in the range post “click” sound stimulation and the peak-to-peak voltage in the range pre “click” sound stimulation. The data processing used Python3 with Pandas, Numpy, Matplotlib, and SciPy libraries.

### Hippocampal brain slice recordings

Time-mated rats were obtained from the Monash Animal Research Platform. The animals were humanely killed on day 18 of pregnancy and the brains were isolated from the fetuses and mounted in ice-cold sucrose- cutting solution in a vibrating microtome (Campden Instrument Integraslice 7550MM, UK), The hippocampal region was cut into 300µm thick slices (blade frequency 80Hz, speed 0.04ms^-1^). All procedures were performed with approval of the Monash University Animal Ethics Committee (project 26803) and followed the National Health and Medical Research Council (NHMRC) of Australia guidelines for the use of animals in research.

rGO-50 electrodes (0.2 mm × 10 mm) were attached to silver wires with silver conductive epoxy and dried at 70 °C for 1 hour. The electrodes were then inserted into glass micropipettes, with the rGO-coated end exposed at the sharp tip and the silver wire at the blunt end. Both ends of the micropipettes were sealed with Araldite epoxy. Brain slices were mounted in a Warner recording chamber on an upright microscope. Each stimulating electrode was placed over the stratum radiatum region of the hippocampus near the rGO-50 electrode. Ag/AgCl electrodes in Glass micropipettes (2–5 MΩ resistance) filled with physiological saline solution were used as a contrast group. Prior to recording with rGO-50 electrodes, Ag/AgCl electrode in a glass micropipette was used as a control to verify the performance of the stimulation electrode and brain slice. Signals were recorded using an Axoclamp-2A amplifier and analyzed with Clampfit 10 software. The Ag/AgCl electrodes have a small diameter (3-10 μm) whereas the graphene-based fibers were 200 μm in diameter. Thus, when stimulated with graphene-based fibers, the current may be spread over a larger area, resulting in a smaller fEPSP amplitude.

## SUPPORTING INFOMATION

Figures S1-S13 and Tables S1-S10

## AUTHOR INFORMATION

The authors declare no competing financial interest.

## ACKNOWLEDGMENT

This work was funded by the NHMRC of the Australian Government (APP1122055), the National Natural Science Foundation of China (No. 5220020670), the Jiangsu Department of Technology Natural Science Fund (BK20220276), and the Dushu Lake Leading Talent Project (MSRIPIF8001006). The Bionics Institute acknowledges the support of the Victorian Government through the Operational Infrastructure Support Program.

