## Supplementary Information for "Graphene-based micro-electrodes with reinforced interfaces and tunable porous structures for improved neural recordings"

† Equal first authors.

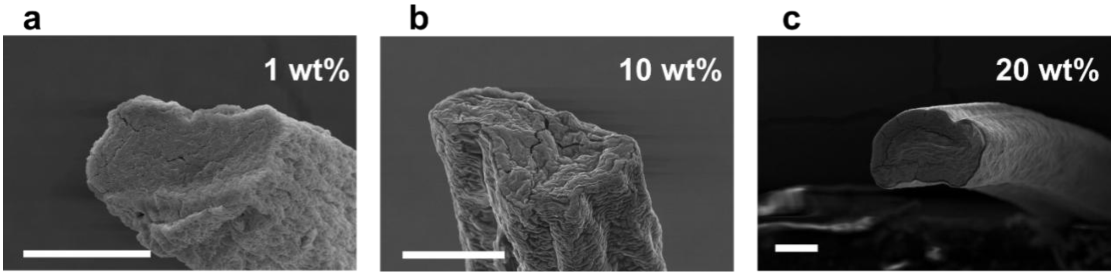


Figure S1. Cross sections of rGO fibers coagulated in baths with different sucrose concentrations and dried at 90 °C. rGO fibers were prepared with a) 1 wt%, b) 10 wt% and c) 20 wt% sucrose in the coagulation bath. Scale bars: 50 µm.

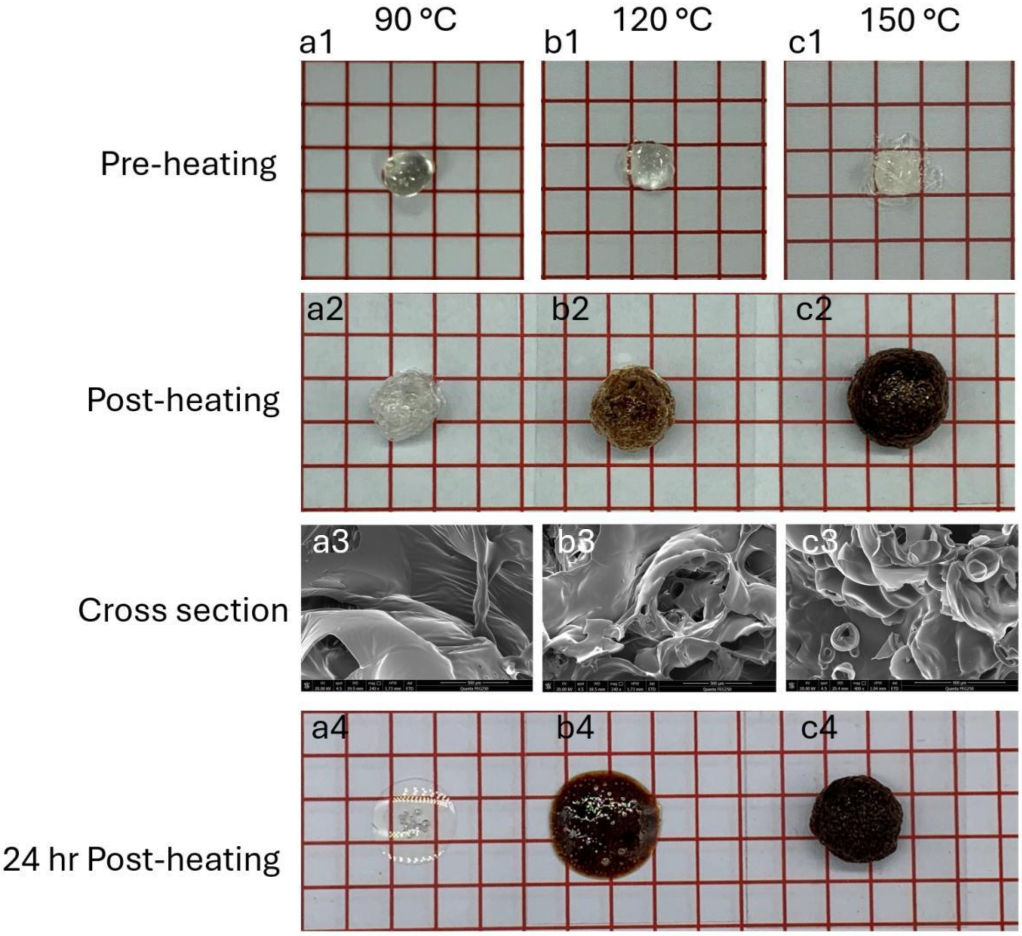


Figure S2. Caramelization of sucrose under vacuum. The concentrated sucrose droplet (figure S2a1) expanded after drying at 90 °C (figure S2a2). The white surface is a clear indicator that no caramelization happens at this temperature. Large bubbles were observed from the cross section of the dried droplet. When the sucrose (figure S2b1) was heated to 120 °C, the droplet exhibited a higher degree of expansion with a color change to light brown (figure S2b2). Cross section was found with additional smaller bubbles (figure S2b3) compared to the droplets dried at 90 °C. These are clear evidence that the caramelization of sucrose starts to happen at 120 °C under vacuum. With further increase of the heat treatment temperature to 150 °C, the sucrose droplet (figure S2c1) expanded to a larger size than the one treated at 120 °C and the color turned into dark brown (figure S2c2), indicating a higher degree of caramelization. More bubbles or ruptured bubbles due to gas release were found (figure S2c3). The expansion of size with a higher degree of caramelization at high temperatures is also reflected in the diameter of rGO-50 fibers (figure 1e). 24-hr post thermal treatment, the droplets in 90 °C (figure S2a4) and 120 °C (figure S2b4) groups melted but one in 150 °C (figure S2c4) maintained its spherical shape, indicating a higher degree of caramelization at 150 °C.

Scale bars: the side of each red square is 5.0 mm. Before the heating process, each concentrated sucrose droplet was around 5 x 5 mm. After heating under vacuum at 90, 120, and 150 °C, it is noted that the colors of heated sucrose at higher temperatures became darker, indicating a higher degree of caramelization. Meanwhile, the expansion in volume supports the gas release. Apart from large bubbles formed from water vaporization at 90 °C, voids formed from smaller bubbles were also observed from the cross sections. The generation of pores are likely attributed to both water vaporization and sucrose caramelization. Additionally, 24 hr after heating, sucrose heated to 90 and 120 °C remelted while sucrose heated to 150 °C maintained a sphere. This phenomenon was caused by the higher degree of caramelization at 150 °C.

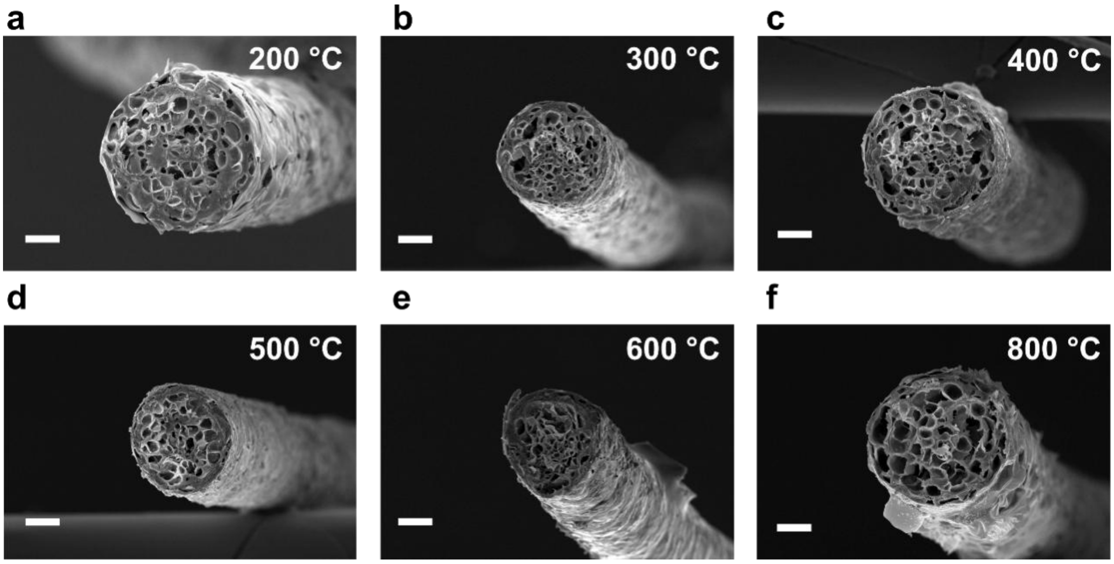


Figure S3. Cross sections of rGO fibers coagulated in a bath with 50 wt% sucrose and reduced at different temperatures. GO fibers were kept at a) 200 °C, b) 300 °C, c) 400 °C, d) 500 °C, e) 600 °C and f) 800 °C for 1 hr during the thermal reduction process. Scale bars: 50 μm.

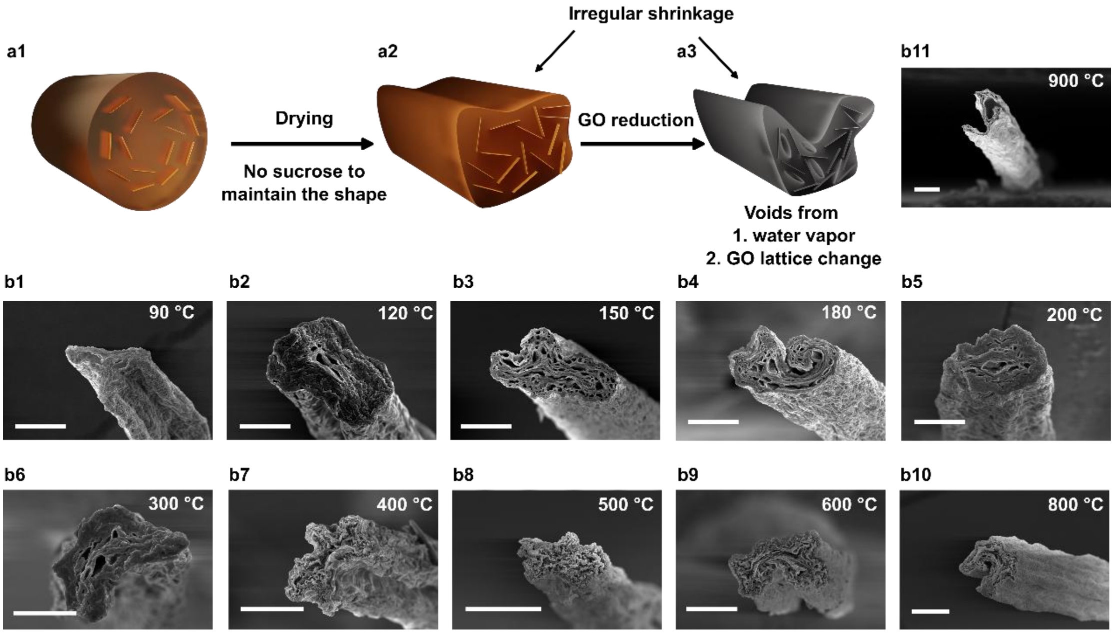


Figure S4. Mechanism of formation of porous rGO fibers without sucrose. a1) GO hydrogel fibers are prepared by the coagulation of GO flakes in a rotating bath with 10 wt% CaCl2 aqueous solution. The fibers are then transferred to a tube furnace for thermal reduction under vacuum. a2) Without the existence of sucrose, the removal of water during the drying process leads to irregular shrinkage of the fiber. a3) GO reduction at higher temperatures intensified the shrinkage of the fiber. Voids form from water vaporization and GO lattice change. b1-b11) Cross sections of fibers reduced at 90, 120, 150, 180, 200, 300, 400, 500, 600, 800, 900 °C, respectively. Scale bars: b1-b11) 50 μm.

Coagulated GO dispersion in $CaCl_2$ was in the state of a hydrogel (figure S4a1). Without sucrose, the removal of water caused shrinkage of fiber in the initial drying process (figures S4a2, S4b1). Above 120 °C, the vaporization of water and GO lattice shrinkage during the reduction process created voids in the GO fiber and intensified deformation (figures S4a3, S4b2-b11).

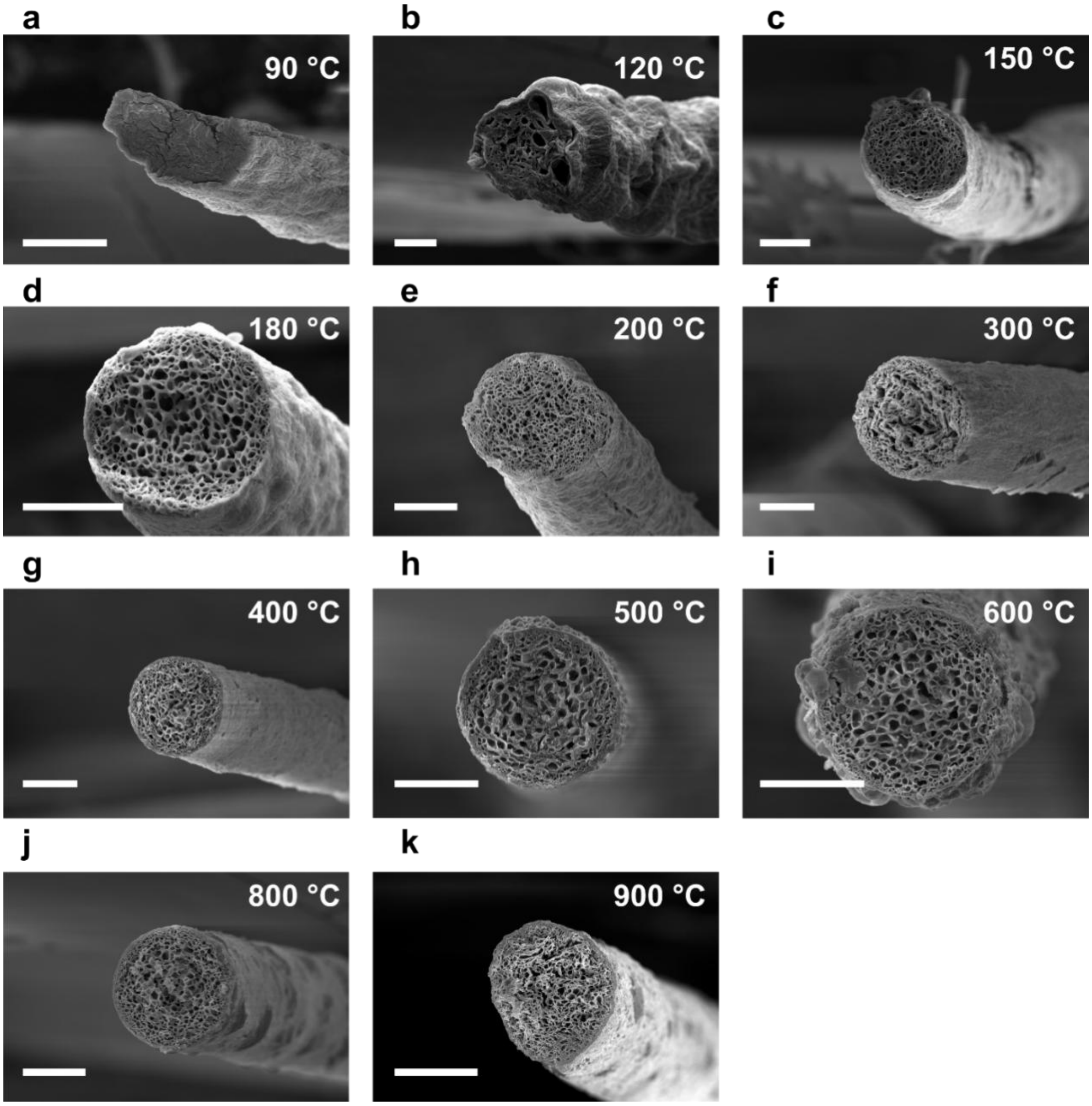


Figure S5. Cross sections of rGO fibers coagulated in a bath with 5 wt% sucrose and reduced at different temperatures. GO fibers were kept at a) 90 °C, b) 120 °C, c) 150 °C, d) 180 °C, e) 200 °C, f) 300 °C, g) 400 °C, h) 500 °C, i) 600 °C, j) 800 °C or k) 900 °C for 1 hr during the thermal reduction process. Scale bars: 50 µm.

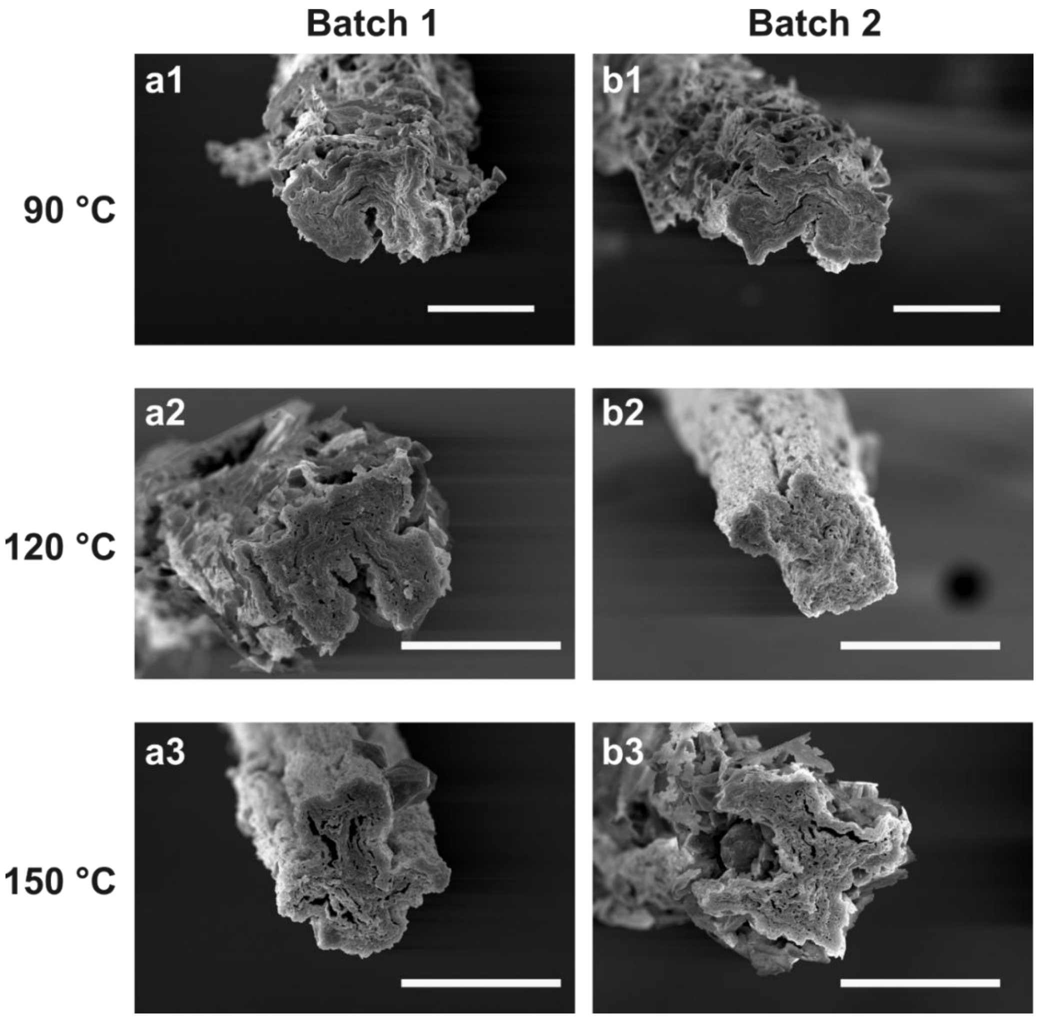


Figure S6. Cross sections of rGO fibers coagulated in 5 wt% sucrose + 10 wt% $CaCl_2$ bath and then soaked in 10 wt% $CaCl_2$ to remove sucrose in the fibers. a) and b) two independent batches of samples were prepared and evaluated. a1) and b1) fibers reduced at 90 °C; a2) and b2) fibers reduced at 120 °C; a3) and b3) fibers reduced at 150 °C. Scale bars: 50 µm.

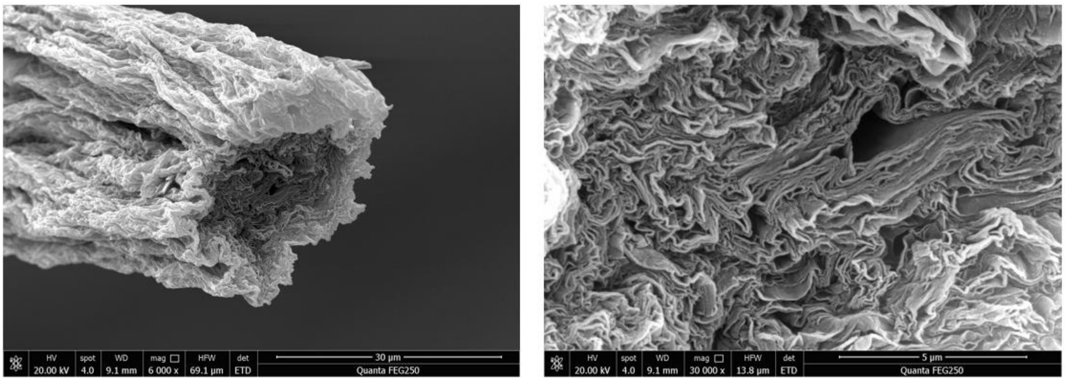


Figure S7. Cross sections of HI reduced rGO fibers. The GO fiber was coagulated in 5 wt% sucrose + 10 wt% $CaCl_2$. The fiber was reduced using HI vapor at 85 °C for 14 hr. The fiber was rinsed in water and ethanol for a few cycles to remove iodine and sucrose.

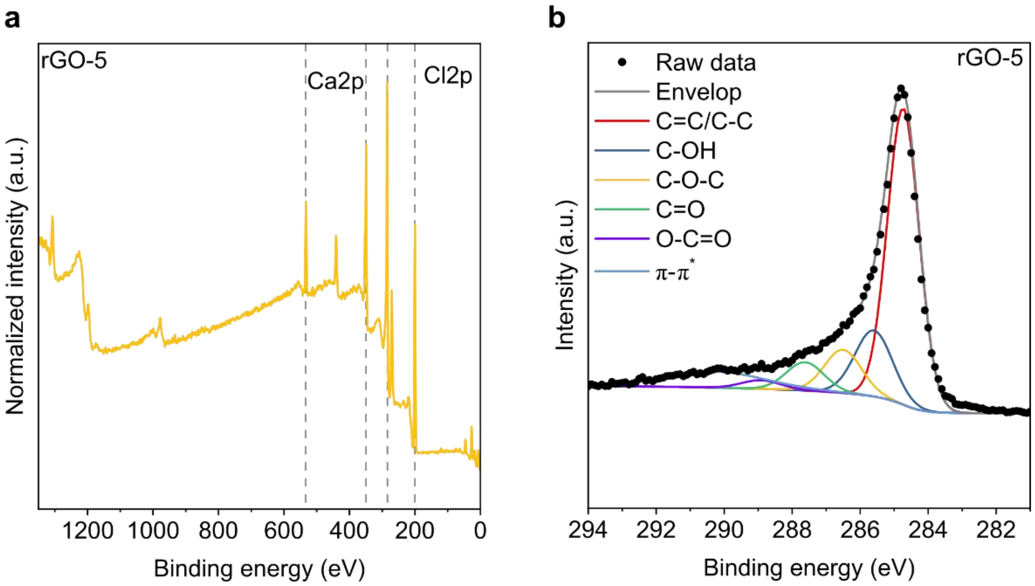


Figure S8. XPS of rGO-5. a) XPS survey and b) C1s spectra of rGO-5.

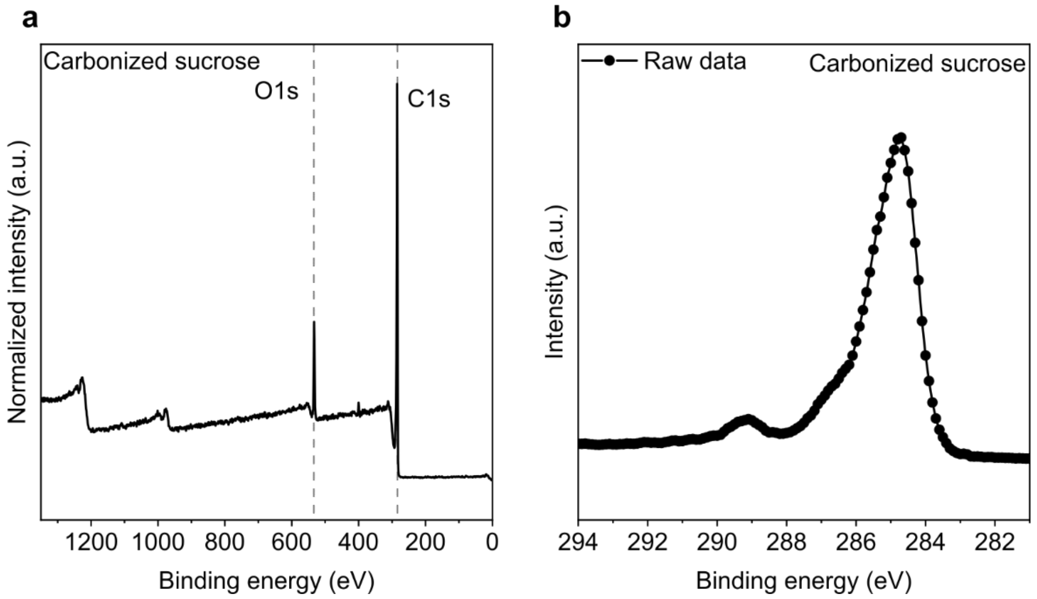


Figure S9. XPS of carbonized sucrose. a) XPS survey and b) C1s spectra of carbonized sucrose. Pristine sucrose crystals were heated to 900 °C under vacuum.

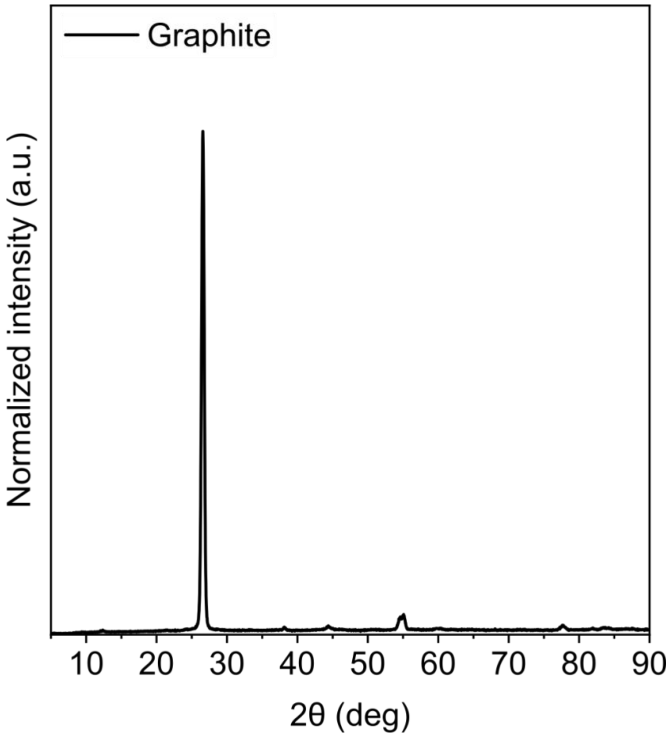

Figure S10. XRD spectra of graphite.

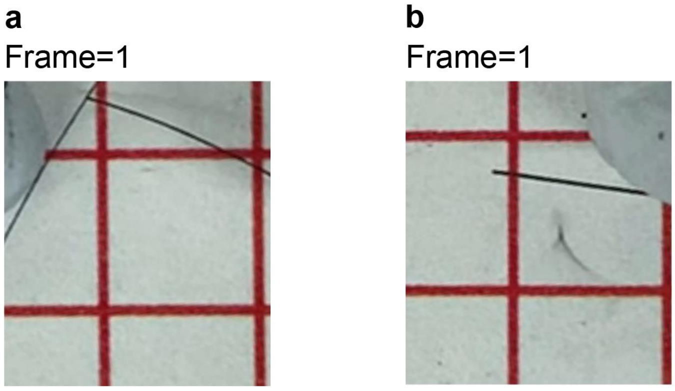


Figure S11. The next frame of a) rGO-0 and b) rGO-5 showing their breakage. The video was recorded at 30 fps. Scale bars: b1, b2) the side of each red square is 5.0 mm.

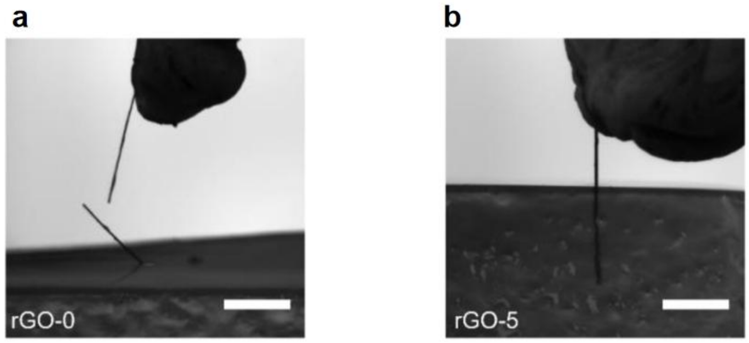


Figure S12. Insertion test of d1) An rGO-0 fiber bent to breakage during the insertion process. d2) An rGO-5 fiber successfully inserted into the gel.

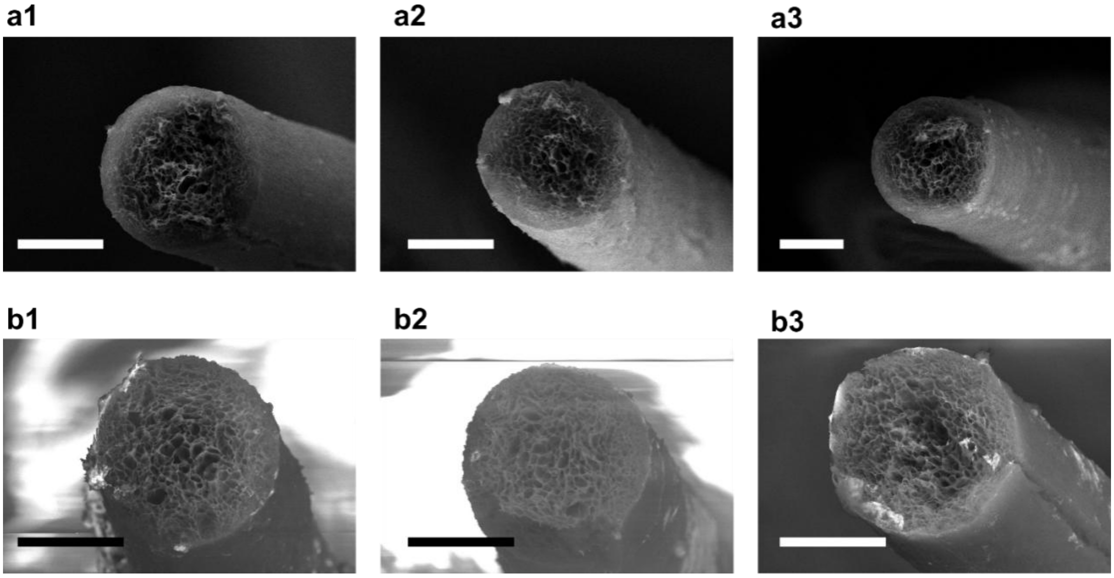

Figure S13. Cross sections of rGO-5 a1) - a3) pre-aging and b1) - b3) post-aging. Scale bars: 50 μm.

Table S1. A summary of porosity and fiber diameter of rGO-0, rGO-1, rGO-5, rGO-10, rGO-20, rGO-30 and rGO-50. Mean±SEM.

| | Porosity | n number | Fiber diameter (μm) | n number |
|---|---|---|---|---|
| rGO-0 | 72.7±1.5 | 5 | 103.9±3.9 | 5 |
| rGO-1 | 88.7±1.8 | 5 | 93.8±2.3 | 5 |
| rGO-5 | 85.4±1.5 | 5 | 130.1±4.6 | 9 |
| rGO-10 | 77.2±0.2 | 5 | 137.4±3.0 | 5 |
| rGO-20 | 78.4±1.5 | 5 | 124.2±2.7 | 7 |
| rGO-30 | 75.2±2.4 | 5 | 152.1±4.9 | 6 |
| rGO-50 | 77.7±0.4 | 5 | 186.0±11.9 | 4 |

Table S2. Fitting result of C1s XPS spectra for GO.

| GO | Binding Energy (eV) | Ratio (%) | FWHM (eV) |
|---|---|---|---|
| C=C/C-C | 284.45 | 44.28 | 1.48 |
| C-OH | 285.47 | 0.67 | 1.34 |
| C-O-C | 286.51 | 46.67 | 1.32 |
| C=O | 288.02 | 6.02 | 1.34 |
| O-C=O | 288.91 | 2.35 | 1.40 |

Table S3. Fitting result of C1s XPS spectra for rGO-0.

| rGO-0 | Binding Energy (eV) | Ratio (%) | FWHM (eV) |
|---|---|---|---|
| C=C/C-C | 284.78 | 74.89 | 1.10 |
| C-OH | 285.80 | 12.45 | 1.34 |
| C-O-C | 286.52 | 7.50 | 1.32 |
| C=O | 287.66 | 4.23 | 1.34 |
| O-C=O | 288.93 | 0.93 | 1.40 |
| π-π* | 290.37 | n/a | 3.03 |

Table S4. Fitting result of C1s XPS spectra for rGO-5.

| rGO-5 | Binding Energy (eV) | Ratio (%) | FWHM (eV) |
|---|---|---|---|
| C=C/C-C | 284.74 | 63.09 | 1.13 |
| C-OH | 285.60 | 16.71 | 1.34 |
| C-O-C | 286.51 | 10.76 | 1.32 |
| C=O | 287.62 | 7.16 | 1.34 |
| O-C=O | 288.91 | 2.29 | 1.40 |
| π-π* | 290.24 | n/a | 3.50 |

Table S5. A summary of magnitude of impedance at 1 kHz, CSC, CIL of Pt, rGO-0, rGO-1, rGO-5, rGO-10, rGO-20, rGO-30, and rGO-50. Mean±SEM.

| | \|Z\| at 1 kHz (Ω $cm^2$) | CSC (mC $cm^{-2}$) | CIL (μC $cm^{-2}$) | n number |
|---|---|---|---|---|
| Pt | 7.65±0.38 | 1.68±0.07 | 45.20±9.33 | \|Z\|: 5, CSC: 5, CIL: 5 |
| rGO-0 | 1.57±0.36 | 151.03±15.12 | 255.00±33.67 | \|Z\|: 7, CSC: 7, CIL: 5 |
| rGO-1 | 0.49±0.04 | 259.19±23.23 | 432.12±56.29 | \|Z\|: 8, CSC: 6, CIL: 6 |
| rGO-5 | 0.57±0.04 | 404.55±31.01 | 877.90±48.98 | \|Z\|: 7, CSC: 5, CIL: 5 |
| rGO-10 | 0.61±0.05 | 169.33±35.33 | 193.97±26.34 | \|Z\|: 6, CSC: 6, CIL: 5 |
| rGO-20 | 1.13±0.02 | 35.58±3.88 | 115.44±8.01 | \|Z\|: 5, CSC: 7, CIL: 5 |
| rGO-30 | 1.20±0.22 | 19.14±3.61 | 109.74±5.70 | \|Z\|: 6, CSC: 5, CIL: 5 |
| rGO-50 | 2.67±0.24 | 2.14±0.26 | 48.12±8.70 | \|Z\|: 5, CSC: 7, CIL: 5 |

Table S6. A summary of the magnitude of conductivity of rGO-0, rGO-1, rGO-5, rGO-10, rGO-20, rGO-30, and rGO-50. Mean±SEM.

| | Conductivity ($\times 10^3$ S $m^{-1}$) | n number | Turkey post-hoc means comparison (vs rGO-0) |
|---|---|---|---|
| rGO-0 | 5.83±0.10 | 5 | n/a |
| rGO-1 | 12.27±0.41 | 5 | $p < 10^{-6}$ |
| rGO-5 | 12.37±0.49 | 5 | $p < 10^{-7}$ |
| rGO-10 | 10.53±0.44 | 5 | $p < 10^{-4}$ |
| rGO-20 | 12.58±0.77 | 5 | $p < 10^{-7}$ |
| rGO-30 | 10.87±0.45 | 5 | $p < 10^{-5}$ |
| rGO-50 | 12.40±0.83 | 5 | $p < 10^{-7}$ |

**Table S7. A summary of tensile Young's modulus of rGO-0, rGO-1, rGO-5, rGO-10, rGO-20, rGO-30 and rGO-50. Mean±SEM.**

| | Young's modulus (MPa) | n number | Turkey post-hoc means comparison (*vs* rGO-0) |
|---|---|---|---|
| rGO-0 | 229.4±31 | 5 | n/a |
| rGO-1 | 980.0±112 | 4 | $p < 10^{-5}$ |
| rGO-5 | 1510.0±119 | 5 | $p < 10^{-7}$ |
| rGO-10 | 852.0±78 | 4 | $p < 10^{-4}$ |
| rGO-20 | 852.8±31 | 4 | $p < 10^{-4}$ |
| rGO-50 | 140.3±23 | 6 | n.s. |

Table S8. A summary of the ultimate tensile modulus of rGO-0, rGO-1, rGO-5, rGO-10, rGO-20, rGO-30, and rGO-50. Mean±SEM.

| | Young's modulus (MPa) | n number |
|---|---|---|
| rGO-0 | 2.6±0.3 | 5 |
| rGO-1 | 7.1±0.8 | 6 |
| rGO-5 | 18.3±2.9 | 5 |
| rGO-10 | 10.7±3.6 | 4 |
| rGO-20 | 8.0±2.5 | 5 |
| rGO-50 | 2.3±0.5 | 6 |

Table S9. A summary of bending radius of rGO-0, rGO-1, rGO-5, rGO-10, rGO-20, rGO-30 and rGO-50. Mean±SEM.

| | Bending radius (mm) | n number |
|---|---|---|
| rGO-0 | 5.0±0.5 | 11 |
| rGO-1 | 5.3±0.3 | 6 |
| rGO-5 | 13.3±0.8 | 5 |
| rGO-10 | 12.0±0.5 | 6 |
| rGO-20 | 12.5±0.4 | 5 |

Table S10. A summary of SNRs of Pt and rGO-0, rGO-1, rGO-5, rGO-10, rGO-20, rGO-30, and rGO-50 recorded from guinea pig auditory cortex at SPL 60, 80, 100 dB. Mean±SEM.

| | SPL 60 dB | SPL 80 dB | SPL 100 dB |
|---|---|---|---|
| Pt | 6.2±1.0, n=10 | 10.1±1.1, n=10 | 11.5±1.2, n=10 |
| rGO-0 | 8.4±0.7, n=12 | 12.0±0.8, n=12 | 14.1±0.8, n=12 |
| rGO-1 | 9.6±0.6, n=12 | 14.9±0.6, n=12 | 16.7±0.6, n=12 |
| rGO-5 | 9.4±1.0, n=9 | 14.6±1.1, n=9 | 15.3±0.9, n=9 |
| rGO-20 | 7.8±0.5, n=13 | 13.4±1.2, n=13 | 13.9±1.1, n=13 |
